# Spoken Recall Reveals Lexical and Mnemonic Function Differences in Temporal Lobe Epilepsy Patients

**DOI:** 10.64898/2026.08.20.746093

**Authors:** Ayelet M. Rosenberg, Eden Tefera, Zehui Gu, Helen Borges, Aaqib Mansoor, Tanay Shah, Gianna Capozzi, William B. Barr, Simon Henin, Stephen B. Johnson, Anli Liu

## Abstract

**Background and Objectives:** Word-finding difficulty is common in healthy aging and in neurologic disorders, including temporal lobe epilepsy (TLE) and early Alzheimer disease. Standard language measures have limited sensitivity for detecting subtle or longitudinal changes in spontaneous speech. We examined whether natural language processing and acoustic analysis of spoken biographical recall could identify lexical and temporal speech features associated with language and memory performance in TLE.

**Methods:** We conducted a cross-sectional observational study of spoken recall during a Famous Faces biographical memory task. Adults with TLE and healthy controls (HCs) viewed 20 famous faces and spontaneously recalled biographical details. Speech was transcribed and diarized using automated tools. Lexical measures included word counts and lexical index (ratio of rare to common words). Acoustic measures included utterance and pause duration and pause frequency. Features were compared between groups and correlated with neuropsychological measures, including Montreal Cognitive Assessment (MoCA), Boston Naming Test (BNT), delayed recall, education, and biographical recall accuracy.

**Results:** Eighty-one adults participated (51 TLE, 30 HCs). Lexical measures did not differ between groups. In TLE, lexical index correlated with BNT performance (r□=0.62) and MoCA score (r□=0.35). Compared with HCs, participants with TLE produced shorter utterances (5.54 ±2.50 vs. 6.59 ±2.63; Cohen’s d=0.41, 95% CI -0.05 to 0.86, p=0.041), shorter pauses (0.61 ±0.21 vs 0.67 ±0.22, Cohen’s d=0.29, 95% CI -0.16 to 0.74, p=0.043), and more frequent pauses (10.81 ±3.30 vs 9.29 ±3.59, Cohen’s d=-0.45, 95% CI -0.90 to 0.01, p=0.036). Higher education was associated with longer utterances, longer pauses, and lower pause frequency, without evidence of a diagnosis-by-education interaction. Faster utterance rate was associated with better biographical recall in both groups, while higher pause rate was associated with worse recall in TLE.

**Discussion:** Speech-derived lexical and temporal features from naturalistic recall capture clinically relevant variation in language and memory-related performance. Although lexical output did not distinguish TLE from HCs, lexical richness tracked naming and global cognition in TLE, while temporal speech features related to recall performance. These findings support the potential of automated speech analysis as a digital behavioral biomarker for word-finding difficulty in neurologic populations.

## INTRODUCTION

Word-finding difficulty (WFD) is a common cognitive complaint in aging^1^ and in neurologic disorders, including Alzheimer disease and temporal lobe epilepsy (TLE).^2,3^ Clinically, WFD is difficult to localize and quantify because it can reflect disruption at multiple levels of the speech-language network, including lexical retrieval, semantic and episodic memory, executive control, and speech production.^3^ Standard clinical measures of WFD, including confrontation naming and semantic or phonemic fluency tasks, remain valuable but have important limitations.^4–6^ They are labor-intensive, produce relatively sparse behavioral data, and may be insensitive to subtle abnormalities in naturalistic speech, especially in individuals with high cognitive reserve.^7,8^ As a result, clinically meaningful changes in everyday language may be missed.

Precise tools for speech assessment are crucially needed in patient cohorts where language dysfunction may be subtle or sporadic. In Alzheimer’s disease, naming and word-finding problems can precede dementia and predict subsequent decline.^9^ In TLE, particularly dominant TLE, patients may produce less informative speech, pause more often, and rely on filler words despite preserved conversational fluency.^2^ Such deficits may arise from dysfunction of left posterior-basal temporal language regions,^2^ or from transient disruption by interictal epileptiform discharges (IEDs) in medial temporal and lateral temporal neocortical networks.^10– 12^

Automated analysis of naturalistic speech potentially represents a quantitative, precise, and scalable approach to this problem. Computational linguistic and acoustic methods can quantify lexical choice and temporal speech structure with greater resolution than conventional bedside testing. In TLE, quantitative studies of naming have demonstrated reduced accuracy and slower response times during object and famous face naming.^13^ In neurodegenerative disease, speech-based biomarkers can distinguish cognitively normal adults from those with mild cognitive impairment and Alzheimer dementia.^14–16^ Temporal measures, such as pause rate, appear particularly promising and have been associated with underlying amyloid and tau burden even in cognitively normal individuals.^9^

We recently found that patients with left TLE showed impaired biographical recall for famous faces compared with healthy controls and patients with right TLE.^17^ This finding was subsequently replicated using semi-automated methods.^18–21^ Here, we extend that work by analyzing recorded spoken recall collected from the Famous Faces task to derive quantitative measures of verbal fluency, defined as lexical and temporal features of spontaneous speech. We hypothesized that lexical features would index vocabulary richness, whereas temporal speech features would reflect mnemonic efficiency during recall. We further tested whether these speech-derived metrics relate to established measures of global cognition, naming, fluency, and memory, including the MoCA,^22^ Boston Naming Test (BNT), Rey Auditory Verbal Learning Test (RVLT), and biographical recall accuracy.

## METHODS

### Subjects

We analyzed audio recordings from 81 adults: 51 with temporal lobe epilepsy (TLE) and 30 healthy controls (HCs), collected at NYU Langone Health from 2018-2024. Recruitment was described previously.^17^ The study included all eligible participants with temporal lobe epilepsy and healthy controls for whom complete speech recordings were available. No formal sample size calculation was performed because this was a retrospective analysis of an existing dataset. Participants were aged 18-60 years. HCs were required to have a Montreal Cognitive Assessment (MoCA)^22^ score ≥26/30 and no self-reported neurologic, psychiatric, or sleep disorder; psychoactive medication use; or alcohol/recreational drug use within 24 hours of testing. TLE participants were required to have a MoCA score ≥ 22/30. A subset of TLE participants also underwent clinical neuropsychological testing, including the Boston Naming Test (BNT), F-A-S fluency, and Rey Auditory Verbal Learning Test (RVLT).

### Standard Protocol Approvals, Registrations, and Patient Consents

All study activities were approved by the NYU Institutional Review Board. All HCs and patients with TLE consented to participate in tasks and share relevant portions of their medical records.

### Famous Faces Task

As previously described,^17^ the Famous Faces (FF) task, adapted from the Iowa Famous Faces Test,^23^ assessed remote memory through face recognition and free biographical recall. Participants viewed 20 famous faces from politics, sports, and entertainment active between 2008 and 2017.^24^ If a face was recognized, participants were asked to recall as much biographical information as possible. To reduce bias from unequal cultural exposure, analyses were restricted to celebrities recognized by each participant. Testing was conducted either in person or remotely via secure WebEx during the COVID-19 pandemic. Audio responses were recorded and stored on HIPAA-compliant local servers. A keyword dictionary for each celebrity was generated from a randomly selected half of HC responses.

### Speech Transcription and Diarization

Audio from the FF task was transcribed with Whisper X (large-v2)^25^ and diarized with pyannote.audio (version 3.1).^26–28^ Speaker labels were assigned using interview structure and known interviewer prompts. WhisperX employs a method of weakly supervised speech recognition trained on 680,000 hours of labeled audio. This model has been found to be relatively comparable to human transcription accuracy and outperforms previously used supervised ASR models.^25^ Pyannote.audio is an open-source Python toolkit for speaker diarization, meaning it can identify and distinguish between different speakers in an audio recording. Pyannote combines various neural building blocks into a simple diarization pipeline, which minimizes diarization error rate.^29^ This speaker diarization method has previously proven effective in diarizing free-flowing natural conversation of patients with autism.^30^ Human reviewers corrected speaker-labeling errors but not transcription errors. Diarization error rate, calculated from false alarms, missed detections, and speaker confusion relative to ground truth duration,^27^ was low (mean 4.0% ± 8.3%).

### Speech Analysis and scoring

Transcripts were analyzed in Python using spaCy, as previously described.^17^ SpaCy processes unstructured text and returns structured data with extensive linguistic information. SpaCy was used to calculate word counts and lexical index (**Figure 1A,1D**). Total word count was defined as all transcript tokens; content word count included nouns, verbs, adverbs, and adjectives (**Figure 1B**). Lexical index was defined as the ratio of low- to high-frequency words. The spaCy package, wordfreq, was used to measure the Zipf frequency of each word.^31^ The wordfreq package measures word frequency based on seven large and diverse word databases such as SUBTLEXus and OpenSubtitles, which collect data from television subtitles,^32,33^ Twitter, Wikipedia, etc. These databases have been shown to better reflect contemporary language and correlate more strongly with performance in word processing tasks.^32^ The Zipf frequency of a word is the base-10 logarithm of the number of times it appears per billion words. Low-frequency words were identified as those with a Zipf frequency of ≤3, or less than once per million words. Words with a Zipf frequency of 4 or more, appearing at least once in ten thousand times, were considered high frequency. We omitted words with a Zipf frequency of 0, because these are not present in the vocabulary of the wordfreq package and are almost always misspellings. We also omitted words that were marked as proper nouns.

**Figure 1.**
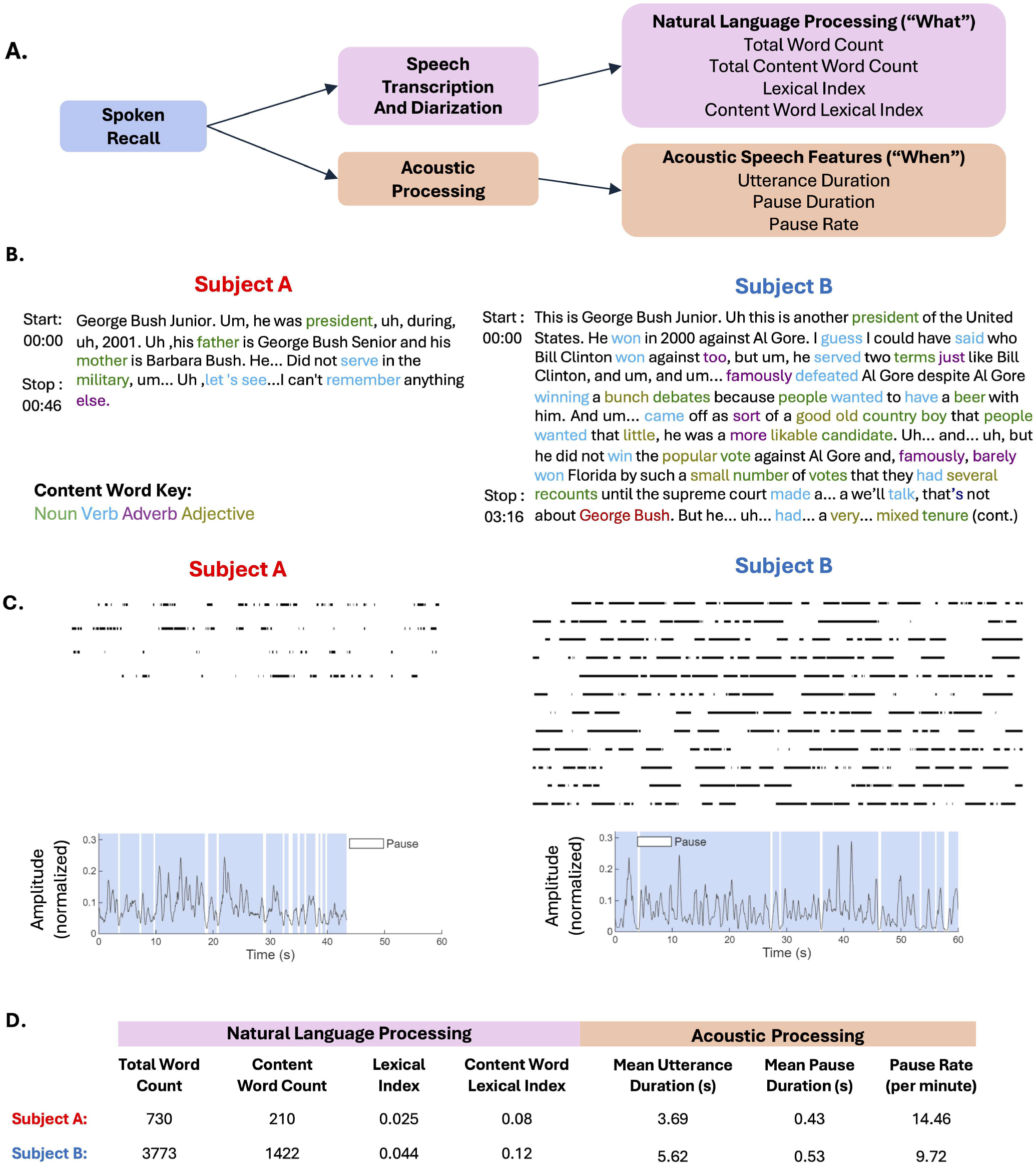
Processing pipeline for extraction of linguistic and acoustic speech features from the Famous Faces (FF) task. **A**. Spoken recall from the Famous Faces (FF) task was transcribed through automated (Whisper Al) means,then speaker identity was assigned via pyannote and validated by humans. The transcript was analyzed via NLP to examine the content of speech, including lexical index and content words. Spoken recall was also analyzed for ratebased features of speech, including mean utterance duration, mean pause duration, and pause rate. **B**. Example speech transcription from two patients with temporal lobe epilepsy, with content word categories (nouns, verbs, adverbs, adjectives) color coded. **C**. Representation of acoustic processing of the same two speech segments shown in B. Visualization of extraction of pause and utterance features. **D**. NLP and acoustic speech feature metrics across the full FF task for the two example patients.

### Acoustic Processing

Audio recordings were processed in MATLAB using custom scripts to derive acoustic speech features (**Figure 1A, 1C**). Only the participant’s speech was analyzed based on the diarized transcripts. Recordings were converted to mono when necessary and resampled to a sampling rate of 16 kHz. Signals were amplitude-normalized by dividing the maximum absolute amplitude so that values ranged between –1 and 1. A smoothed amplitude envelope was then derived from the absolute value of the waveform using a moving average filter. An adaptive threshold was then applied to classify each time point as speech or silence. Silence segments <250ms were reclassified as speech,^34^ and speech segments <150ms as silence, to reduce noise-related misclassification (**Figure 1C**). Contiguous speech and pause segments were then identified. Mean utterance duration and mean pause duration were calculated in seconds, while pause rate was calculated as pauses per minute of speech (**Figure 1D**).

### Scoring and Statistical Analysis

Recall transcriptions were scored for the percent of accurate details included in the keyword dictionary. Generated by two blinded reviewers (T.S. and G.C.). Continuous measures were calculated for each group (TLE, HC), including age, total MoCA score, MoCA phonetic fluency score, MoCA delayed recall score, and FF Task Recall performance. These were reported as means and standard deviations. Categorical variables, such as sex, handedness, and educational level, were reported as counts and percentages. Continuous demographic and cognitive performance was compared between TLE and HCs using independent-samples t-tests, whereas Pearson’s chi-square tests of contingency were performed on categorical data. A subset of TLE patients also had available neuropsychological testing (Boston Naming Test, F-A-S phonemic fluency, and RVLT delayed recall, see summary **eTable 2**), for which we report the means and standard deviations.

For continuous metrics derived from subject transcripts such as word counts, content word counts, lexical index, content word lexical index, and acoustic analysis such as utterance duration, pause duration, and rates, the Shapiro-Wilk test was used to test normality of distribution. Because metrics were not all normally distributed, we used nonparametric tests for all analyses. The Wilcoxon Two-Sample Test was used for comparing derived continuous metrics between TLE and HC groups. Relationships between continuous variables were assessed using Spearman’s nonparametric correlations. Associations between education level and speech metrics were assessed using Spearman rank-order correlations. Linear regression models were used to test interactions between independent variables, which we defined as the fixed variables (e.g., Diagnosis, MoCA, BNT. Phonetic fluency, etc.) and dependent variables, which we defined as the spoken-recall derived lexical and temporal metrics.

### Data Availability

The anonymized patient data and related clinical trial study documents will not be publicly shared as they are being used for development of a clinical trial. Data will be made available after publication of this article and up to 5 years after publication. The authors will share the data with qualified investigators whose proposal of data use has been approved by the principal investigator (A.L.).

## RESULTS

### Subjects

Eighty-one (81) patients (51 TLE, 30 HCs) were included in this retrospective cross-sectional analysis (**Table 1**). There were no group-level differences in sex (63% female, p = 0.60) or education status (78% college or above, p = 0.24). TLE patients were older than HCs, with mean ages of 33.2 + 9.6 years (TLE) and 27.9 + 8.5 years (HC) (Cohen’s d = -0.57, 95% CI -1.04 to -0.11, p = 0.014). Fewer TLE patients than HCs were right-handed (78% vs. 97%, p = 0.037). TLE patients scored lower than HCs on the MoCA (26.6 + 2.3 vs. 28.9 + 1.1, Cohen’s d = 1.17, 95% CI 0.68 to 1.68, p<0.0001) and MoCA delayed recall (3.5 + 1.4 vs. 4.6 + 0.7, Cohen’s d = 0.96, 95% CI 0.48 to 1.44, p <0.0001), which is an artifact of the lower MoCA eligibility criteria for TLE patients. TLEs scored lower than HCs on the FF Task Recall (33.8 + 11.8 vs. 38.8 + 11.7, Cohen’s d = 0.42, 95% CI -0.04 to 0.88, p = 0.04). There were no differences in MoCA phonetic fluency performance (F words in a minute) (TLE: 14.2 + 4.1, HC: 15.3 + 4.3, Cohen’s d = 0.25, 95% CI -0.21 to 0.70, p = 0.30).

**Table 1.**
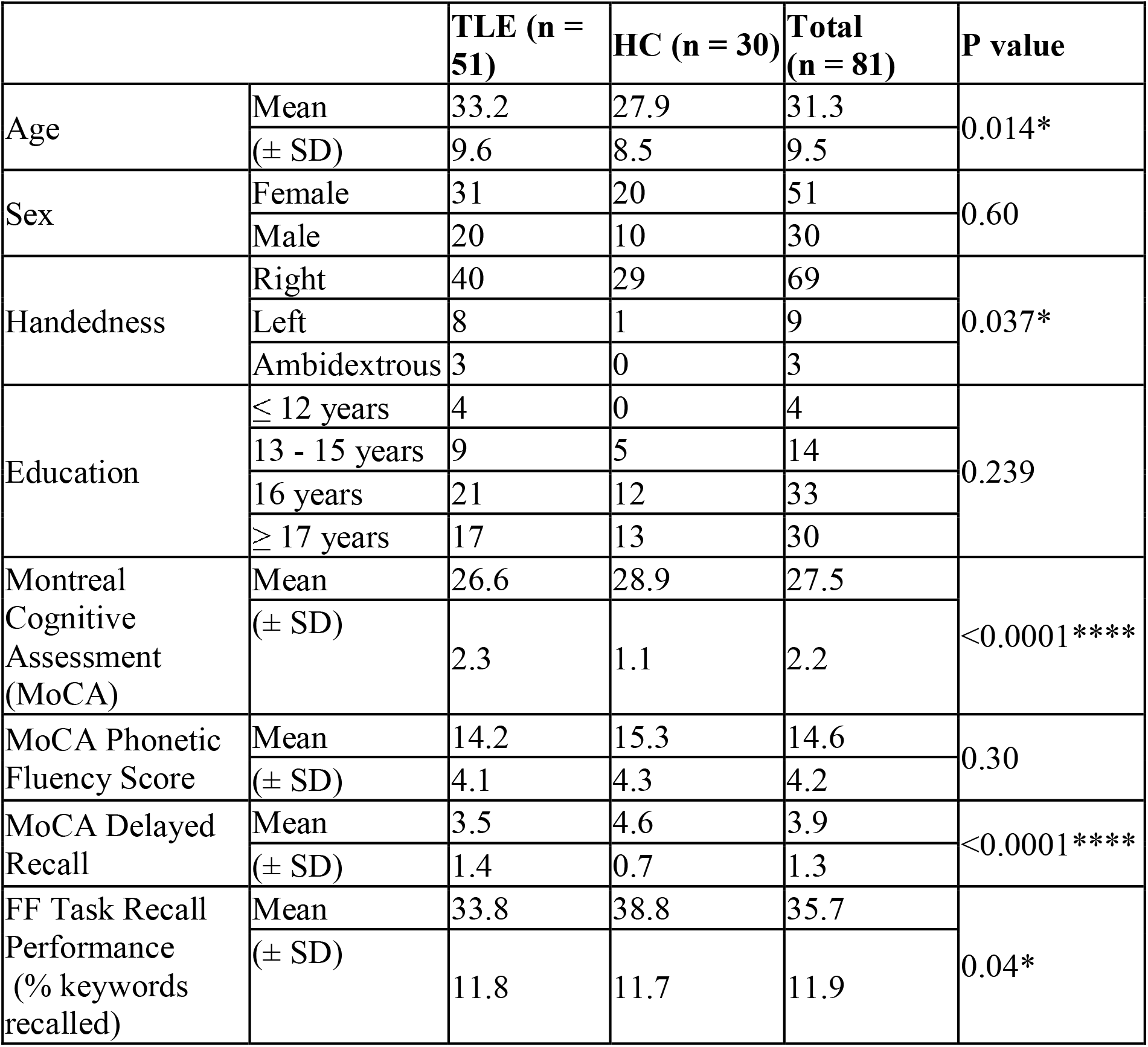
Subject Demographics. Participant demographics and cognitive characteristics by group. Continuous demographic and neuropsychological variables were compared between TLE and healthy control (HC) participants using independent-sample t-tests. Categorical variables were compared using Pearson’s chi-square tests. MoCA total score out of 30. MoCA Phonetic Fluency Score was the number of F words generated in a minute.MoCA delayed recall is out of 5. TLE = temporal lobe epilepsy, HC = healthy controls. Note: * indicates p < 0.05, and **** indicates p < 0.0001.

LTLE (26) and RTLE (25) patients were similar in age, sex, handedness, education, MoCA scores, MoCA Phonetic Fluency score, and FF Recall. However, MoCA delayed recall scores were lower for LTLE patients compared to RTLE patients (3.0 ± 1.6 words vs. 3.9 ± 1.1 words out of 5 words total, Cohen’s d = - 0.69, 95% CI -1.26 to -0.12, p = 0.019) (**eTable 1**). In addition, among TLE patients who had clinical neuropsychological testing, LTLE and RTLE patients performed similarly on the BNT (LTLE: N = 15, 50.3 ± 8.3 words vs. RTLE: N = 18, 51.3 ± 6.5 words out of 60 total words, Cohen’s d = -0.13, 95% CI -0.81 to 0.56, p = 0.88); Phonetic Fluency F-A-S (LTLE: N = 15, 35.5 ± 11.1 vs. RTLE: N = 16, 41.3 ± 16.2 words, Cohen’s d = -0.41, 95% CI -1.13 to 0.30, p = 0.24); and RVLT (LTLE: N = 11, 8.1 ± 3.0 words vs. RTLE: N = 12, 10.1 ± 2.8 words out of 15 total, Cohen’s d = -0.66, 95% CI -1.40 to 0.07, p = 0.08) (**eTable 2**).

Missing data were minimal. One participant had missing age data and was excluded from analyses involving age. One participant had a missing Montreal Cognitive Assessment (MoCA) score and was excluded from analyses involving MoCA. All other analyses included participants with available data only.

### Lexical features were similar, but speech rates differed between HCs and TLE patients

There were no group-level differences between TLE and HC subjects for Total Word Count (965.1 ± 875.9 vs. 1015.2 ± 863.4; Cohen’s d: 0.058, 95% CI -0.39 to 0.51, p = 0.68), Content Word Count (347.0 ± 327.9 vs. 372.9 ± 328.2; Cohen’s d: 0.079, 95% CI -0.37 to 0.53, p = 0.75), Lexical Index (LI) (0.028 ± 0.011 vs. 0.029 ± 0.012; Cohen’s d: 0.15, 95% CI -0.30 to 0.60, p = 0.63), and Content-Word LI (0.074 ± 0.030 vs. 0.079 ± 0.034; Cohen’s d: -0.14, 95% CI -0.59 to 0.31, p = 0.61). (**Table 2**). Using NLP methods to identify parts of speech, we examined whether the number of specific content words, such as nouns and pronouns, differed between TLE and HC patients, but they did not. However, TLEs and HCs differed across all temporal speech features, with TLEs having shorter mean utterance durations compared to HCs (5.54 ± 2.50 secs vs. 6.59 ± 2.63 secs; Cohen’s d = 0.41, 95% -0.05 to 0.86, p = 0.041), shorter mean pause durations (0.61 ± 0.21 sec vs 0.67 ± 0.22 sec, Cohen’s d = 0.29, 95% CI -0.16 to 0.74, p = 0.043), and more frequent pauses per minute (mean 10.81 ± 3.30/ min vs. mean 9.29 ± 3.59/min, Cohen’s d = -0.45, 95% CI -0.90 to 0.01, p = 0.040) (**Table 2, Figure 2A**). Of note, these rate-based features were also related to educational level, with less-educated subjects speaking in shorter utterances (r□ = 0.24, p = 0.028), shorter pauses (r□ = 0.31, p = 0.0043), and a higher pause rate (r□ = -0.26, p = 0.021) (**Table 2, Figure 2B**). However, there was no interaction between diagnosis (TLE vs. HC) and educational level for any of the metrics (p = 0.057, p = 0.097, p = 0.164, respectively). Together, these findings suggest that TLEs and HCs produced similar speech output and lexicon, but TLE patients and less educated subjects spoke in shorter utterances with shorter and more frequent pauses compared to HCs.

**Table 2.**
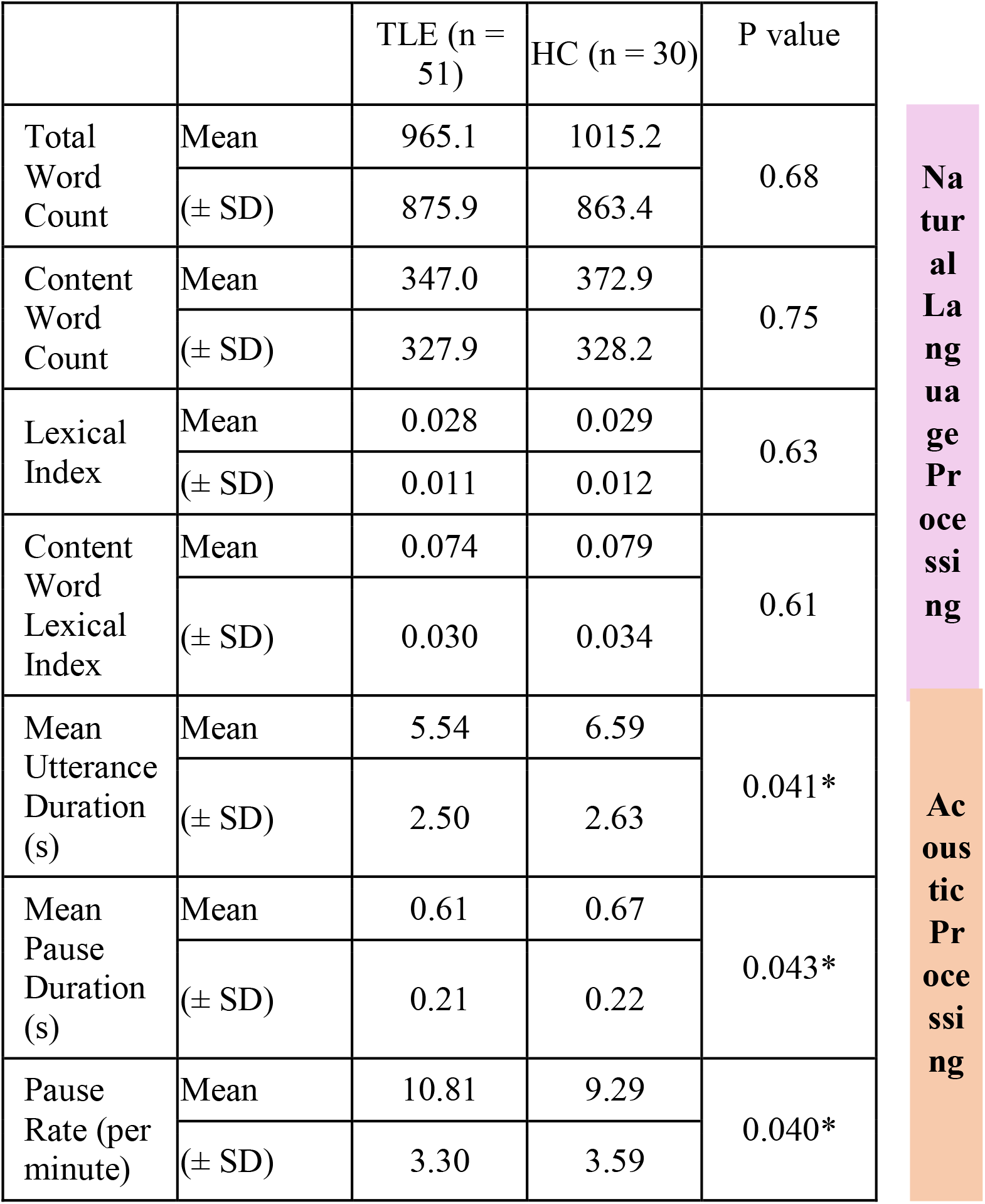
Results. Lexical and Temporal Speech measures between TLE patients and healthy controls. Lexical features derived from Natural Language Processing (NLP) and Rate-based features of spoken recall derived from the acoustic envelope were compared between TLE and HC participants using nonparametric Wilcoxon for two-sample tests. Total words include all words spoken by subject, including filler words (um, uh, like…). Content word count included nouns, verbs, adjectives and adverbs. Lexical Index is defined as the ratio of low to high frequency words used in spoken recall. HC = healthy control; TLE = temporal lobe epilepsy. Note: * indicates p < 0.05.

**Figure 2.**
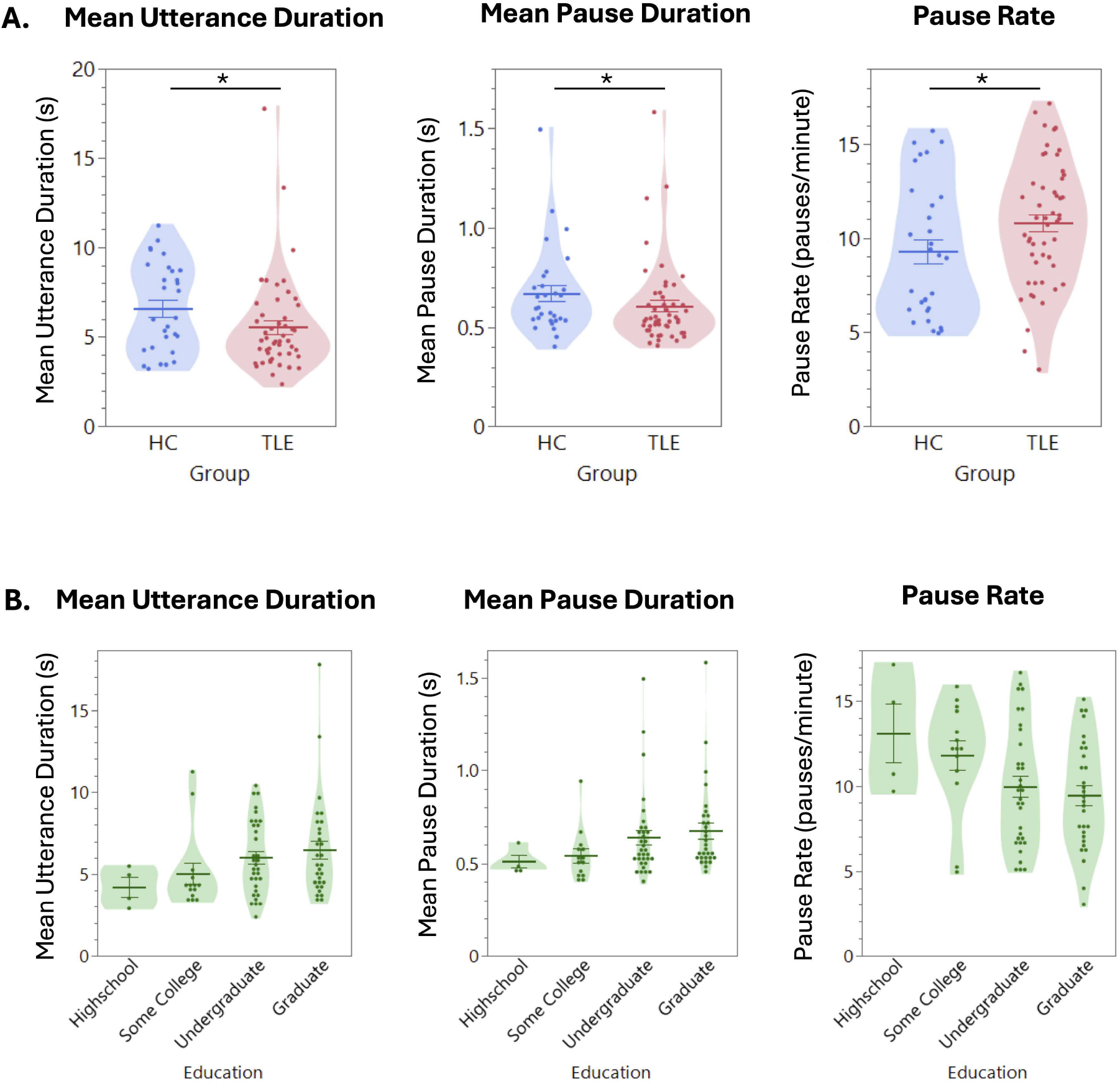
Temporal features of Spoken Recall Differ by Diagnosis and Education Level. **A**. Violin plots comparing mean utterance duration, mean pause duration, and pause rate between TLE patients and HCs. All group differences were statistically significant (Wilcoxon two-sample test). *Indicates p <0.05. **B**. Violin plots showing each acoustic metric across education levels (all participants). Higher education was associated with longer utterances, longer pauses, and lower pause rates, consistent with Spearman correlations (p = 0.24, 0.31, -0.26, respectively). There were no significant group x education interactions for any metric, indicating that the effect of education was consistent across TLE and HC participants.

### Lexical Index correlates with MoCA and Boston Naming Test (BNT)

MoCA correlated strongly with Lexical Index (LI) in TLEs (r = 0.35, p = 0.013) but not significantly in HCs (r = 0.35, p = 0.058) (**Figure 3A**). MoCA also correlated strongly with Content Word LI in both TLEs (r = 0.35, p = 0.012) and HCs (**r** =0.42, p = 0.023) (**eFigure 1A**). BNT performance correlated strongly with LI (r =0.62, p=0.0001, **Figure 3B)** and Content Word LI (r =0.63, p=0.0001) in TLE patients (**eFigure 1B**). To determine whether LI was an indirect measure of specific biographical details, we tested the relationship between specific content words (ratios of nouns, pronouns, and verbs to total words) and amount of biographical detail (as detailed in Tefera et al.^17^) and did not find a significant relationship. We note that LI in our cohort of HCs and TLE patients was normally distributed (p=0.27), whereas MoCA and BNT were highly skewed (p<0.0001, p<0.0006, respectively) and were limited by ceiling effects (**Figure 3C**).

**Figure 3.**
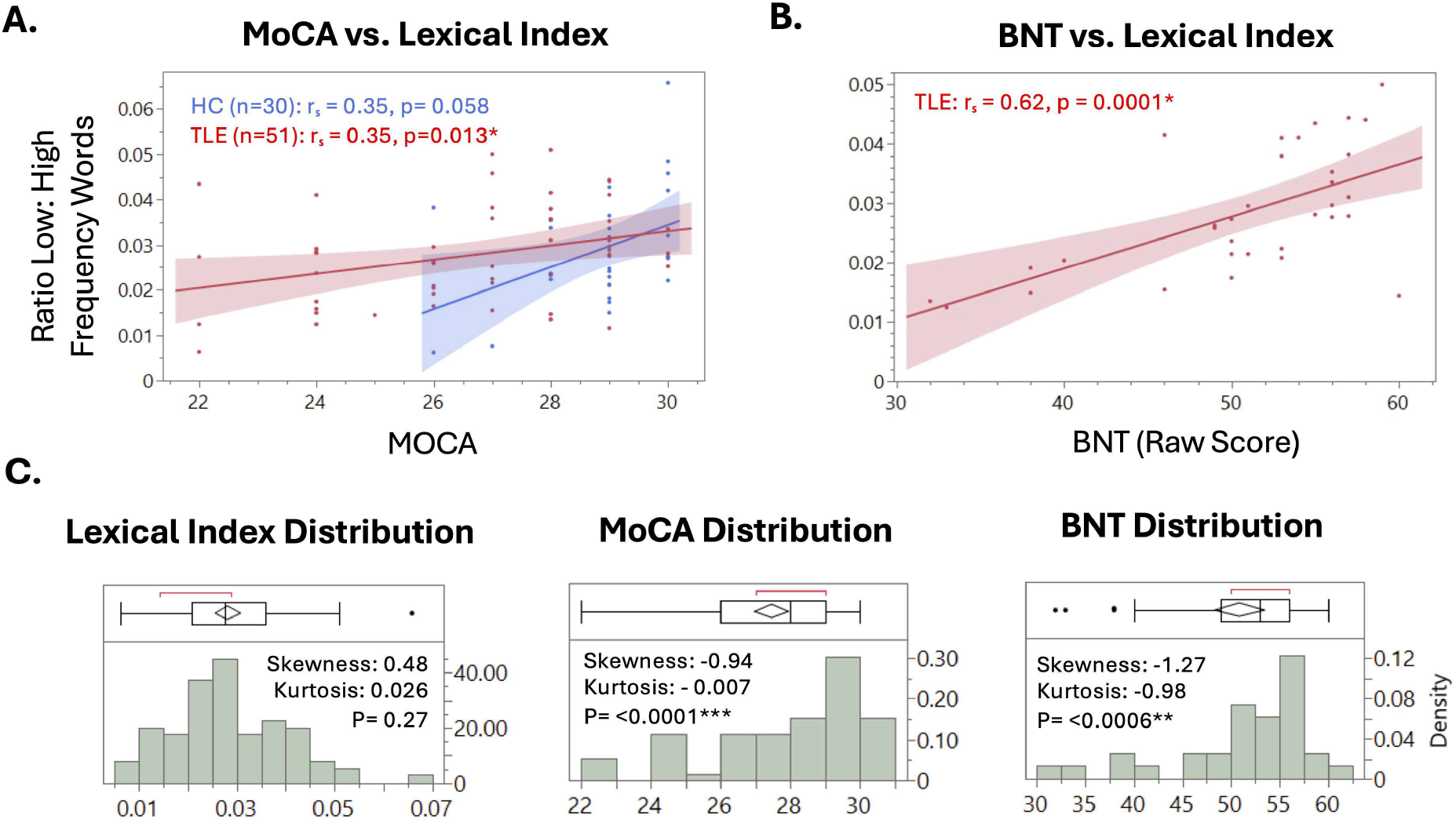
MoCA and BNT correlate with Lexical Index in TLE patients. **A**. Scatterplot showing the relationship between MoCA score and Lexical Index. In TLE patients, a significant correlation was observed (p= 0.35, p = 0.013), whereas no significant association was observed in the HC group (r = 0.35, p = 0.058). B.Scatterplot showing a significant correlation between BNTscore and Lexical Index in TLE patients (r = 0.62, p= 0.0001). Points represent individual participants and are colored by group (TLE in RED, HC in BLUE). BNT was collected in TLE patients only. TLE = temporal lobe epilepsy; HC = healthy control. C. Distribution of Lexical Index, MoCA, and BNTscores. Normality testing indicated that Lexical Index was normally distributed, whereas MoCA and BNTwere significantly skewed and showed evidence of ceiling effects.

To test the specificity of LI to naming –and not memory or executive function – we compared LI to MoCA Phonetic Fluency, MoCA Delayed Recall, and RVLT performance. There was no significant relationship between LI and any of these neuropsychological measurements (MoCA Phonetic Fluency: TLE – r□= 0.27, p = 0.06, HC - r□ = 0.25, p = 0.19; MoCA Delayed Recall: TLE-r□ = 0.24, p = 0.10, HC – r□= 0.28, p = 0.12; RVLT: TLE – r□= -0.13, p = 0.50) (**eFigure 2**). For HCs but not TLE patients, LI was correlated with memory scores on the FFT, as measured by the percentage of keywords remembered for familiar famous faces (**eFigure 4**).^35^

In summary, we did not find group-level differences between TLE and HCs in lexical counts or vocabulary use, suggesting that this TLE patient population has a relatively intact language function. However, even among a high-performing subject pool, we have validated a lexical index of spoken recall against standardized general cognitive (MoCA) and naming tests (BNT). LI appears to be specific to naming, as it did not correlate with standard verbal memory or executive function testing. However, unlike the MoCA and BNT, LI measures a broader range of performance between subjects and is not limited by ceiling effects, a concern among more highly educated patients. Furthermore, LI can be derived from natural speech outside of the clinic and thus used to assess a patient’s lexicon over time.

### Pause rate and other temporal features differed between TLE and HCs but did not correlate with standard memory metrics

Subjects with a TLE diagnosis and with less education spoke in shorter utterances and with shorter and more frequent pauses compared to HCs and subjects with more education, respectively (**Figure 2**). To determine whether pause rate and related temporal features of speech related to established cognitive tests of verbal episodic memory (MoCA delayed recall, RVLT), we tested the relationship between MoCA, MoCA Delayed Recall, and RVLT with Pause Rate. We did not find a relationship between pause rate and MoCA, MoCA Delayed Recall, or RVLT in TLE or HCs (MoCA : TLE-r□ = -0.05, p = 0.72, HC – r□ = 0.20, p=0.29; MoCA Delayed Recall: TLE – r□ = 0.01, p = 0.93, HC - r□ = -0.05, p = 0.78; RVLT: TLE – r□ = -0.04, p = 0.83) (**eFigure 3**). In the case that these memory metrics were not sensitive to subtle memory decline, we compared pause rate to biographical recall and found that they were negatively correlated in TLE participants (r□ = -0.31, p = 0.026), but not in HCs (r□ = -0.34, p = 0.07) (**eFigure 4**). Utterance rate was also negatively correlated with biographical recall in both TLE participants (r□ = -0.36, p = 0.01) and HCs (r□ = -0.41, p = 0.02), suggesting that temporal characteristics of speech may reflect aspects of memory retrieval not captured by traditional cognitive measures (**eFigure 4**).

## DISCUSSION

Word finding difficulty (WFD), or “tip-of-the-tongue” phenomenon, is one of the most common complaints of healthy aging^1,36^, but may also signify risk of later cognitive decline.^37^ WFD is a common complaint in patients with temporal lobe dysfunction and may reflect impairment in speech-language or episodic memory networks. Standard testing has identified cognitive phenotypes in patient cohorts. For example, left (dominant) TLE patients often show deficits in confrontation naming and verbal fluency tasks.^38–40^ Postoperative naming decline is a well-documented consequence of left temporal lobe resections for epilepsy, especially for low-frequency and proper nouns.^41,42^ Alzheimer’s disease (AD) often presents with subtle word-finding and naming deficits, which can predict conversion to dementia in longitudinal studies.^43,44^

However, standard testing may be insensitive to subtle declines. For example, although approximately two-thirds of TLE patients report memory difficulty, standard testing only detects objective deficits in half of these patients. Clinical testing often fails to detect preclinical AD by up to 2 decades.^7,45^

By applying natural language processing and acoustic analysis to spoken biographical recall, we demonstrate that lexical and temporal speech features capture distinct cognitive functions. Vocabulary, indexed by the ratio of low- to high-frequency words, correlates with naming ability (BNT) and global cognition (MoCA). In contrast, pause and utterance are associated with depth of biographical recall, despite being missed by standard metrics of naming, phonemic fluency, or list-learning memory. Spontaneous recall may capture lexical and temporal signals relevant to language and memory function, respectively.

These speech metrics can capture clinically meaningful information. TLE patients spoke in shorter phrases and more frequent pauses compared to HCs; as did less educated patients. These group-level measurements mirrored differences in biographical recall performance,^17^ suggesting a sensitive biomarker for memory function. We have previously demonstrated that naturalistic recall performance on a famous faces task correlates with general cognition (MoCA) and verbal list learning (RVLT), but is not limited by ceiling effects. Our findings suggest that natural speech-derived measures may be more sensitive to subtle naming or memory dysfunction than conventional instruments.

Our findings are consistent with prior work utilizing speech biomarkers to diagnose clinical stages of Alzheimer’s disease. Studies have leveraged a variety of speech tasks, including picture description, story recall, and verbal fluency tasks.^9,46–49^ Slowed speech rates, increased pause frequency, and reduced phonation correlate with disease stage.^50^ Vocabulary becomes less varied, precise, and informative with more advanced disease.^16,14,51^ Individuals with positive amyloid- or tau-AD biomarkers demonstrate subtle but consistent slowing of speech and reduced lexical diversity (i.e., high-frequency, vague words), semantic richness, and narrative coherence, compared to controls. These changes are observed even in the context of normal clinical neuropsychological scores.^51–54^

However, our study extends prior approaches by applying a computational framework to quantify lexical performance. By operationalizing use of low versus high frequency words, we suggest the potential for a highly sensitive metric to track subtle loss of words over time. Additionally, these computational models can be continuously updated to reflect evolving vernacular language patterns, providing an adaptable framework for tracking lexical change over time that is less dependent on static normative comparisons derived from specific English-speaking populations.

Our findings contribute to a growing body of literature supporting the use of speech analysis to provide diagnostic information. Lexical and temporal metrics covary with naming and mnemonic tasks, validating domain-specific information. Because these metrics are not limited by ceiling effects, they represent a powerful new approach to track cognitive function in an individualistic, serial manner. Our findings and others^51–56^ suggest that recall performance may index mnemonic efficiency or retrieval efforts that are missed by standard neuropsychological tests. Moreover, because speech samples may be collected on an ongoing basis, they could potentially capture the rare to occasional lapses in word-finding missed in one-shot clinical assessments, especially in patients with high cognitive reserve.

We acknowledge several limitations of the study. Healthy controls were younger and had higher MoCA scores than the TLE group, which may have influenced group comparisons. We also did not assess mood, subjective cognitive complaints, or other factors that may shape self-perceived WFD. In addition, this was a retrospective, exploratory study in a modest sample, and confidence intervals were not available for all reported associations.

In summary, speech-based biomarkers have shown promise in mild cognitive impairment and Alzheimer disease, where acoustic and linguistic features can distinguish disease stages and may relate to underlying pathology. Our results extend this framework to epilepsy and support spontaneous speech as a scalable digital behavioral marker of cognitive function. Because audio can be collected repeatedly and analyzed semi-automatically, these methods may be well suited for longitudinal monitoring, including detection of subtle decline, treatment effects, or postsurgical changes.

## Funding

NIH R01NS127954 (A.L.), NIH K23NS104252 (A.L.), NYU FACES (A.L.), NYU Department of Neurology (A.L.)

## Acknowledgments

ChatGPT (GPT 5.5) was used for editing purposes under supervision to improve the clarity and readability of portions of the manuscript text.

## Notes

### Competing Interest Statement

The authors have declared no competing interest.

