## Supplemental Figure 1 for "Spoken Recall Reveals Lexical and Mnemonic Function Differences in Temporal Lobe Epilepsy Patients"

**eFigure 1.**

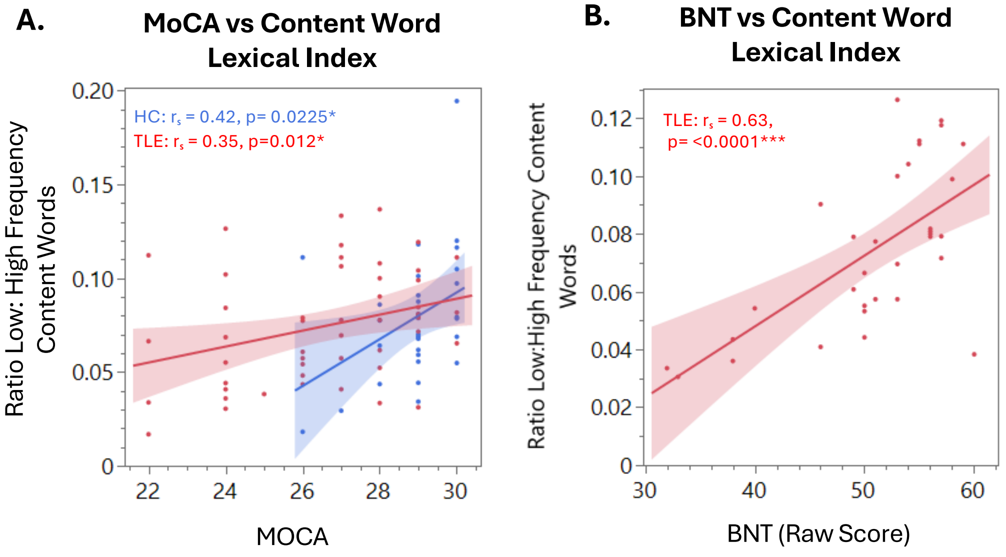


**eFigure 1. Content Word Lexical Index correlates with MoCA and BNT.** Scatterplots showing the relationship between MoCA score and Content Word Lexical Index (**A**) and BNT score and Content Word Lexical Index (**B**). Points represent individual participants and are colored by group (TLE in red, HC in blue). Spearman rank-order correlation coefficients (ρ) are shown. The fitted line represents the overall trend. HC = healthy control; TLE = temporal lobe epilepsy**.**
