## Supplemental Figure 2 for "Spoken Recall Reveals Lexical and Mnemonic Function Differences in Temporal Lobe Epilepsy Patients"

**eFigure 2.**

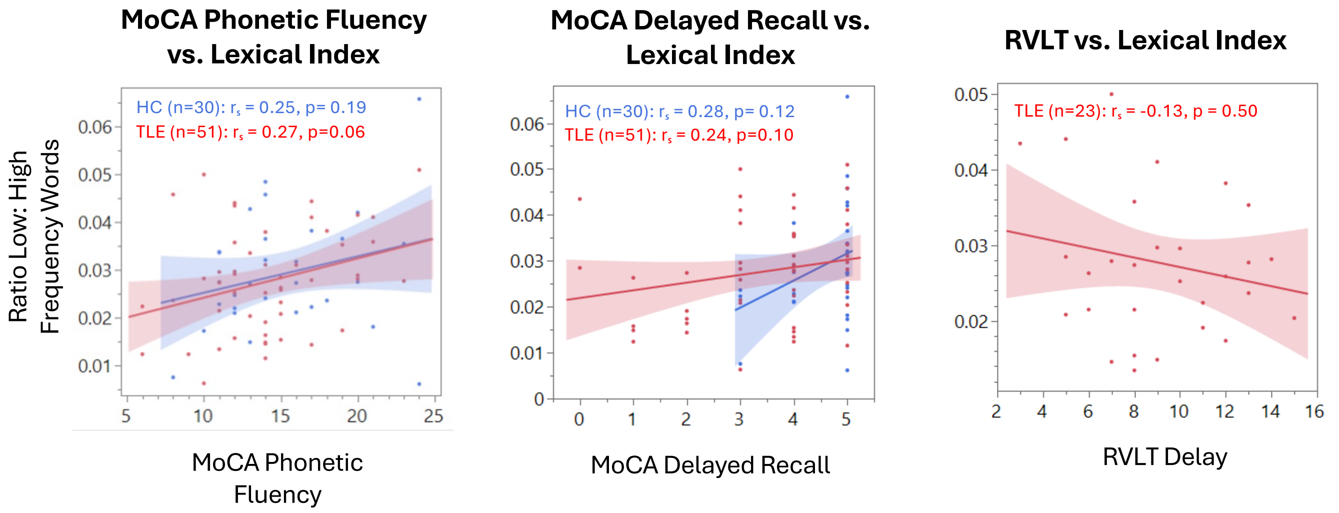


**eFigure 2.** Scatterplots showing the relationship between MoCA Phonetic Fluency (left), MoCA Delayed Recall (middle), and RVLT Delay (right) with Lexical Index. Points represent individual participants. Spearman rank-order correlation coefficients (ρ) are shown. The fitted line represents the overall trend. TLE = temporal lobe epilepsy, HC = Healthy control
