## Supplemental Figure 4 for "Spoken Recall Reveals Lexical and Mnemonic Function Differences in Temporal Lobe Epilepsy Patients"

**eFigure 4.**

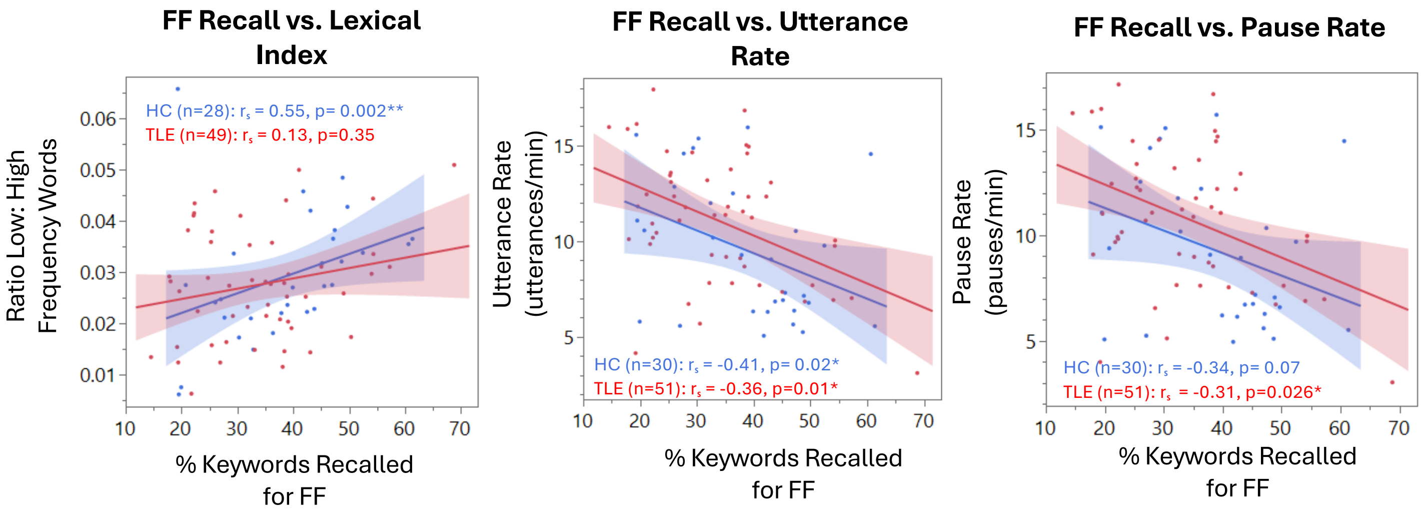


**eFigure 4.** Scatterplots showing the relationship between Lexical Index (left), Utterance Rate (middle), and Pause Rate (right) with Famous Faces Recall scores. Points represent individual participants. Spearman rank-order correlation coefficients (ρ) are shown. The fitted line represents the overall trend. TLE = temporal lobe epilepsy, HC = Healthy control
