## Supplemental Table 1 for "Spoken Recall Reveals Lexical and Mnemonic Function Differences in Temporal Lobe Epilepsy Patients"

**eTable 1. Left vs Right TLE Subject Demographics**

| | | LTLE (n =26) | RTLE (n = 25) | P value |
|---|---|---|---|---|
| Age | Mean | 31.7 | 34.7 | 0.27 |
| | (± SD) | 9.2 | 10.0 | |
| Sex | Female | 15 | 16 | 0.64 |
| | Male | 11 | 9 | |
| Handedness | Right | 21 | 19 | 0.81 |
| | Left | 4 | 4 | |
| | Ambidextrous | 1 | 2 | |
| Education | ≤ 12 years | 3 | 1 | 0.75 |
| | 13 - 15 years | 5 | 4 | |
| | 16 years | 10 | 11 | |
| | ≥ 17 years | 8 | 9 | |
| Montreal Cognitive Assessment (MoCA) | Mean | 26.4 | 26.9 | 0.43 |
| | (± SD) | 2.3 | 2.3 | |
| MoCA Phonetic Fluency Score | Mean | 13.5 | 15.0 | 0.20 |
| | (± SD) | 3.6 | 4.6 | |
| MoCA Delayed Recall | Mean | 3.0 | 3.9 | 0.019* |
| | (± SD) | 1.6 | 1.1 | |
| FF Task Recall Performance (% keywords recalled) | Mean | 33.5 | 34.2 | 0.59 |
| | (± SD) | 11.9 | 11.9 | |

**eTable 1. Participant demographics and cognitive characteristics by TLE Laterality.** Continuous demographic and neuropsychological variables were compared between LTLE and RTLE participants using independent-samples t-tests. Categorical variables were compared using Pearson's chi-square tests. MoCA total score out of 30. MoCA Phonetic Fluency Score was the number of F words generated in a minute. MoCA delayed recall is out of 5. LTLE = Left temporal lobe epilepsy, RTLE = Right temporal lobe epilepsy. Note: * indicates $p < 0.05$
