## Supplemental Table 2 for "Spoken Recall Reveals Lexical and Mnemonic Function Differences in Temporal Lobe Epilepsy Patients"

### eTable 2. TLE only Neuropsychology Testing Scores

| | | | | |
|---|---|---|---|---|
| Neuropsychiatry testing | Boston Naming Test (raw) | N = 31 | Mean | 50.85 |
| | | | (± SD) | 7.29 |
| | F-A-S Phonetic Fluency (raw) | N = 30 | Mean | 38.45 |
| | | | (± SD) | 14.07 |
| | RVLT Delay Recall (raw) | N = 23 | Mean | 9.1 |
| | | | (± SD) | 3.02 |

**eTable 2. Cognitive performance of TLE patients on standard neuropsychological tests.** Mean ± standard deviation (SD) scores are reported for each test. Only TLE patients completed these assessments.
